# Potent LILRB1 D1D2-containing antibodies inhibit RIFIN-mediated immune evasions

**DOI:** 10.1101/2024.08.08.607148

**Authors:** Yizhuo Wang, Hengfang Tang, Wanxue Wang, Ming Li, Chenchen Zhu, Han Dai, Hongxin Zhao, Bo Wu, Junfeng Wang

**Affiliations:** High Magnetic Field Laboratory, Key Laboratory of High Magnetic Field and Ion Beam Physical Biology, Hefei Institutes of Physical Science, Chinese Academy of Sciences, Hefei, Anhui 230031, China; University of Science and Technology of China, Hefei, Anhui 230026, China; Department of Anatomy, School of Basic Medical Sciences, Anhui Medical University, Hefei, China; Hefei China Science Longwood Biological Technology Co., Ltd. Hefei, Anhui 230088, China

**Author notes:** Corresponding authors. (Hongxin Zhao), (Bo Wu), (Junfeng Wang). Those authors contributed equally (Yizhuo Wang, Hengfang Tang).

**Keywords:** *Plasmodium falciparum*, immune evasion, RIFIN, receptor-containing antibody

## Abstract

The spread of drug-resistant malaria parasites presents a major challenge to global efforts in malaria control, increasing the urgency for new treatments and vaccines. A promising approach involves developing antibodies that can counteract the parasite’s immune evasion mechanisms. In this study, we designed a receptor-containing antibody, targeting the D1D2 domain of the LILRB1 receptor, using a structure-based rational approach. We began with the MDB1 antibody as a scaffold and replaced the LILRB1-D3D4 insertion domain with D1D2.v, a high-affinity variant optimized through yeast surface display. The modified D1D2.v-IgG efficiently blocked the interaction between RIFIN#1 (from PF3D7_1254800) and LILRB1, thereby reversing the inhibition of NK cell activity caused by RIFIN#1. To further enhance this effect, we developed NK-biAb, a bispecific antibody based on D1D2.v-IgG that targets both RIFIN#1 and NKG2D receptors. NK-biAb exhibited superior biological performance compared to D1D2.v-IgG alone. These findings provide a clear framework for designing antibodies that target immune evasion in malaria, potentially guiding the development of more effective treatments and vaccines.

## Introduction

Malaria remains one of the deadliest infectious diseases globally, with Plasmodium falciparum being the species most responsible for high mortality rates, particularly in sub-Saharan Africa^1,2^. The emergence and spread of drug-resistant P. falciparum strains pose a significant threat to current malaria control efforts, underscoring the urgent need for new treatments and vaccines^3–5^. The life cycle of Plasmodium falciparum is complex, spanning several developmental stages between the mosquito vector and the human host. Among these, the blood stage is primarily responsible for the clinical symptoms of malaria, including fever and anemia^6,7^. During this stage, P. falciparum invades erythrocytes, where it replicates and displays a variety of variable surface antigens on infected erythrocytes (IEs). These blood-stage parasites are the primary target of acquired immunity. However, by interacting with host immune receptors, these surface antigens allow P. falciparum to evade immune detection, contributing to the slow development of naturally acquired immunity against malaria^7^.

RIFINs, the largest family of surface antigens expressed by P. falciparum, are key players in immune evasion^7,8^. These antigens include numerous isoforms, each capable of binding to different domains of host receptors such as LAIR1 or LILRB1^9–12^ (Fig. 1a). For instance, RIFIN PF3D7_1254800 binds preferentially to the D1D2 domain of LILRB1^9^, while RIFIN PF3D7_1101100 interacts with LAIR1 rather than LILRB1^10^. Blocking these receptor-RIFIN interactions could prevent immune evasion and enhance immune clearance, providing a potential strategy for developing effective malaria treatments and vaccines^13,14^.

Recent studies have identified several receptor-based antibodies containing LAIR1 and LILRB1-D3D4 fragments designed to target these immune evasion mechanisms^15–17^ (Fig. 1b). Unlike conventional antibodies, which bind antigens through their variable regions, these receptor-containing antibodies incorporate LAIR1 or LILRB1 fragments between the variable and constant regions of the heavy chain, enabling them to directly target specific RIFIN antigens. LAIR1-containing antibodies, in particular, have undergone specific mutations that enhance their affinity for RIFINs^16^. In contrast, LILRB1-containing antibodies retain the unmutated LILRB1-D3D4 domain or just the D3 domain in the variable-constant (VH-CH1) elbow region^15^ (Fig. 1c). Although some RIFINs bind to the LILRB1-D1D2 domain, mimicking the interaction of β2-microglobulin with MHC class I molecules, no antibodies targeting this domain have been identified. This may be due to the inherently low affinity between RIFINs and LILRB1-D1D2. Moreover, developing receptor-containing antibodies in vivo is time-consuming, particularly due to the uncontrollable nature of the affinity maturation process.

In this study, we introduce a rational, structure-based approach to design a receptor-containing antibody that targets the D1D2 domain of LILRB1. This newly developed antibody family mimics the natural immune recognition mechanism, providing a novel strategy to competitively inhibit RIFIN-mediated immune evasion. To create a high-affinity D1D2 variant (D1D2.v) capable of antagonizing the interaction between LILRB1 and RIFIN#1 (from PF3D7_1254800), we used yeast surface display to direct the evolution of the LILRB1-D1D2 domain^18–20^. The receptor-containing antibody MDB1 served as the scaffold, and we replaced the LILRB1-D3D4 insertion domain with D1D2.v to generate the D1D2.v-containing antibody, D1D2.v-IgG (Fig. 1d). This newly engineered antibody demonstrated a 110-fold higher affinity for RIFIN#1 than the native LILRB1-D1D2 domain, successfully blocking the interaction and restoring NK cell activity suppressed by RIFIN#1. Building on this success, we further developed a bispecific antibody, NK-biAb, based on D1D2.v-IgG, which includes an NKG2D-specific scFv (KYK2.0) at its C-terminus^21–23^. This design aims to redirect NK cells toward parasite-infected erythrocytes, mimicking CAR-NK therapy^24,25^. NK-biAb showed superior biological activity in enhancing NK cell-mediated cytotoxicity compared to D1D2.v-IgG alone. These findings establish a framework for the development of novel antibodies targeting immune evasion mechanisms, providing new avenues for malaria treatment and vaccine development.

**Figure 1:**
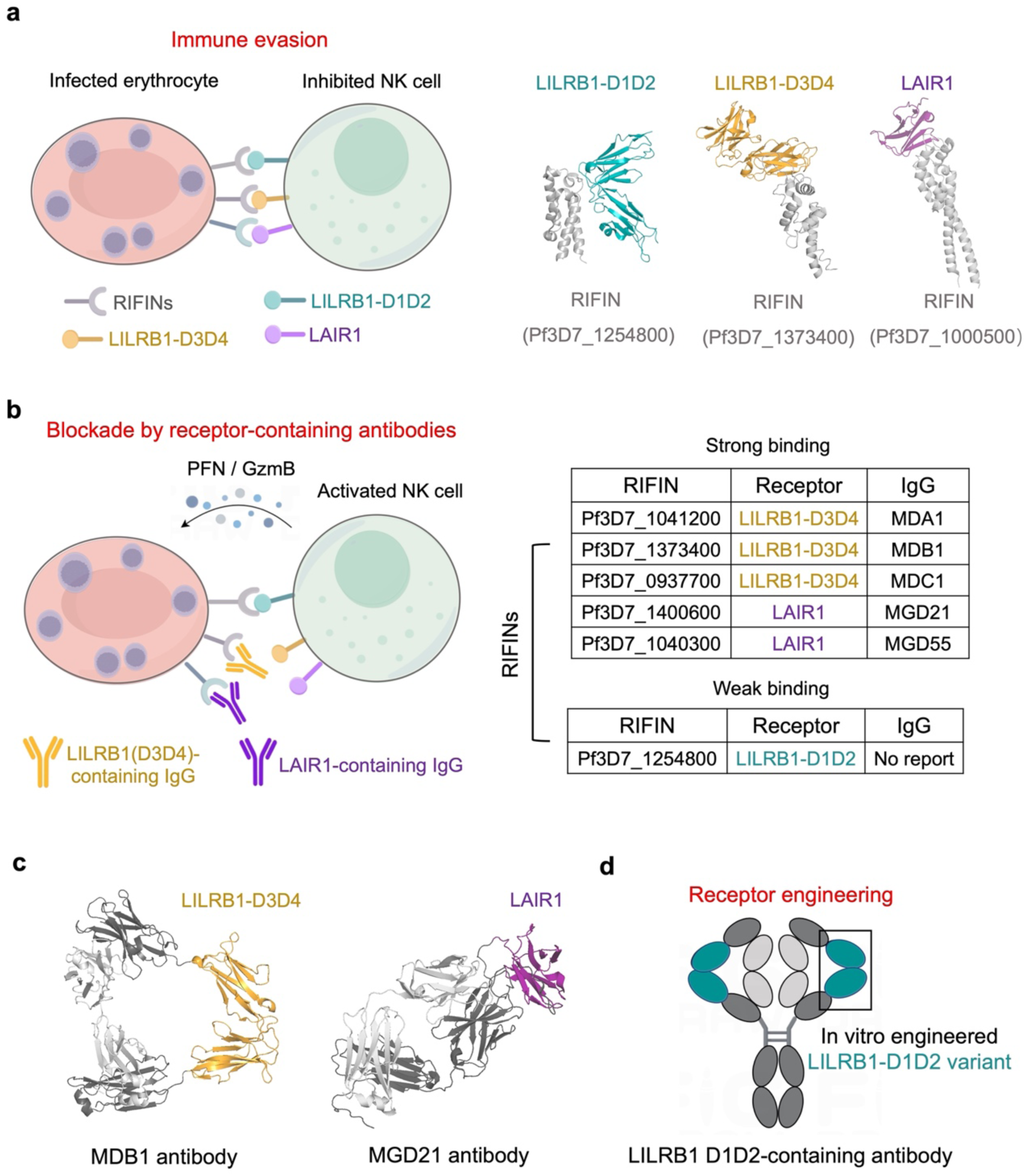
Immune evasion of *P.falciparum* and design of novel receptor-containing antibody. (**a**) Mechanisms of immune evasion of *P. falciparum* and structures of RIFIN and immunoinhibitory receptors. Different RIFIN isoforms bind to their corresponding receptor domains and transduce immunosuppressive signals (PDB: 6ZDX, 7KFK and 7F9N). (**b**) Receptor-containing antibodies in inhibiting immune evasion of *P. falciparum* and identification of LILRB1-and LAIR1-containing antibodies. All reported LILRB1-containing antibodies only have LILRB1-D3D4 domains and no antibodies that contain LILRB1-D1D2 domains were found. (**c**) Two representative receptor-containing antibody. The left one is LILRB1 D3D4-containing antibody MDB1 (PDB:7KHF) and the right one is LAIR1-containing antibody MGD21 (PDB: 7JZ4). (**d**) Strategy for generating high-affinity LILRB1 D1D2-containing antibodies. The affinity-matured LILRB1-D1D2 variant was generated through structure-based affinity maturation in vitro and then the LILRB1-D1D2 variant was grafted into the framework of MDB1 by replacing LILRB1-D3D4 domains.

## Results

### Structure-guided engineering of a high affinity LILRB1 D1D2 variant

RIFIN#1 binds to the extracellular domain (ECD) of LILRB1 through its variable region with a moderate affinity (K_d_ =570 ± 130 nM). LILRB1-ECD comprises four immunoglobulin-like domains (D1-D4), and its complex structure with RIFIN#1 suggests that the interaction is mediated by the D1D2 domain^9^. In order to obtain a high-affinity variant capable of competitively antagonizing LILRB1-RIFIN#1 interaction, we performed structure-guided directed evolution of the LILRB1-D1D2 domain by yeast surface display. A library of LILRB1 D1D2 mutations was designed using the structure of the LILRB1-RIFIN#1 complex as a guide. As shown in Figure 2a, ten LILRB1 D1D2 residues (K42, E68, G97, A98, Y99, I100, Q125, V126, F128, E184) interfacing with RIFIN#1 were altered based on the structural information by randomized PCR. The resulting D1D2 mutant library contained 2x10^7^ transformants. The library was then displayed on the yeast surface, stained with biotinylated RIFIN, and screened to isolate high-affinity binders (Fig. 2b and Fig. S1). In total, eight round of sorting were performed (Fig. S2). Three rounds of sorting with decreasing concentrations of biotinylated RIFIN tetramers (2 μM, 1 μM and 0.3 μM) (Fig. S3) and three rounds of sorting with decreasing concentrations of biotinylated RIFIN monomers (0.3 μM, 0.1μM and 0.02 μM) (Fig. S4) were performed for binder enrichment. For kinetic sorting, enriched library was screened using 0.02 μM biotinylated RIFIN monomers in the presence of approximately a 50-fold excess unlabeled competition RIFIN, with a duration of 2 h and 4 h respectively (Fig. S5). After completion of all eight rounds of sorting, yeast cells were transplanted to obtain individual clones. DNA sequencing results indicate that a single D1D2 mutant clone (named D1D2.v) dominates this enrichment process, and this consensus variant contains a total of nine mutations: K42M, G97S, A98T, Y99F, I100T, Q125E, V126I, F128Q, E184H (Fig. 2c and Fig. S6).

To gain insight into the mechanisms of D1D2.v affinity enhancement, we obtained a structural model of D1D2.v binding to RIFIN#1 using SWISS-MODEL (Fig. 2d). Comparative analysis with the crystal structure of the LILRB1-RIFIN#1 complex show that D1D2.v binds to RIFIN#1 in essentially the same manner as its wild-type counterpart; however, some differences are observed at the binding interface. As shown in Figure 2d, the Q125E mutation establishes a new salt bridge with R233 on RIFIN#1. In addition, the G97S, A98T, and I100T mutations appears to enhance the strength of their H-bonds with RIFIN#1 based on analysis of the H-bond distance.

**Figure 2:**
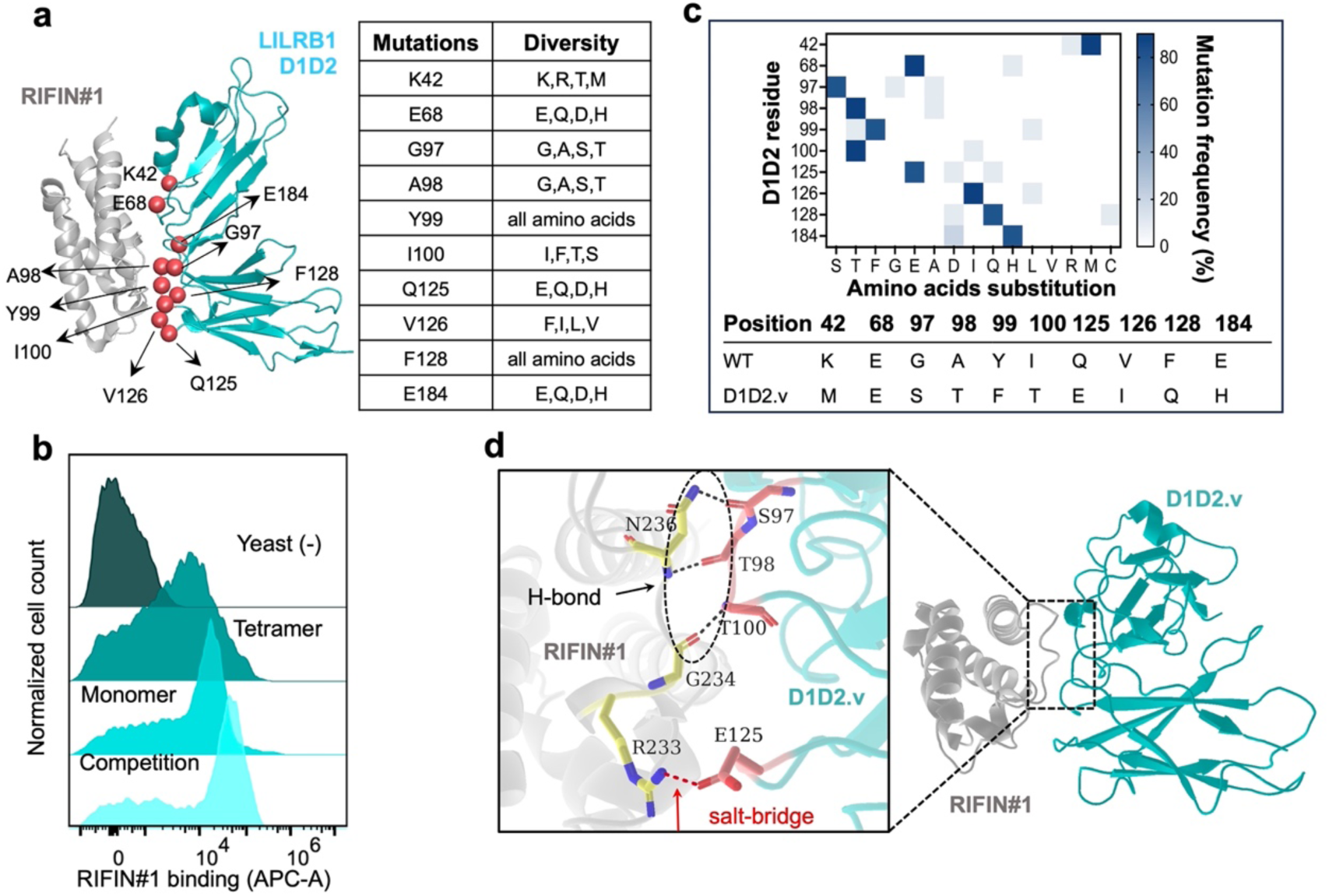
Yeast display and affinity maturation of D1D2 domain. (**a**) (Left) LILRB1 models based on LILRB1-RIFIN#1 complex structure (PDB:6ZDX), where RIFIN#1 is in orange color, LILRB1-D1D2 is in cyan color, and the randomized positions are depicted by red balls; (Right) Table of random positions of the LILRB1-D1D2 interface library with possible amino acid variants. (**b**) Flow cytometry results for the yeast stained with biotinylated RIFIN#1 in tetramer, monomer and competition selection strategy. (**c**) Mutation frequencies (Above) and consensus D1D2.v sequence (Below). (**d**) Structural modeling and interface analysis. The sequences of D1D2.v and RIFIN#1 were imported into the SWISS-MODEL server for initial model generation based on the coordinates of PDB code. The G97S, A98T, I100T and Q125E mutations in D1D2.v increases H-bonding strength targeting RIFIN#1, while a salt-bridge forms between E125 on D1D2.v and R233 on RIFIN#1.

### Design and biochemical characterization of a D1D2.v-containing antibody that antagonizes RIFIN#1-LILRB1 interaction

Recently, several natural LILRB1-containing monoclonal antibodies were identified in malaria-infected individuals, such as MDB1, in which the LILRB1 D3D4 structural domain was inserted into its variable-conatant(VH-CH1) elbow region^15,16^. We then replaced the LILRB1-D3D4 insertion domain in MDB1 with D1D2.v to generate a D1D2-containing antibody, named D1D2.v-IgG (Fig. 3a). To express as the secreted protein, the codon-optimized antibody sequence was genetically linked to a N-terminal Trp2 signal peptide sequence within a pTT5 expression vector. The recombinant antibody was transiently expressed in Expi293F cells and purified from culture supernatant by protein A affinity chromatography. The purity was confirmed by SDS-PAGE analysis under both non-reducing and reducing conditions. To evaluate the RIFIN#1-binding activity of this recombinant antibody, we generated the Fab fragment by enzymatically removing the Fc tag and detected its binding with RIFIN#1 by size-exclusion chromatography (SEC) assay. The results showed that Fab bound strongly to RIFIN#1, forming a tight complex (Figure 3b). Next, we quantified the binding affinity of D1D2.v-IgG to RIFIN#1 using a protein-based ELISA assay with purified LILRB1 ECD as a control. As shown in Fig. 3c, the dissociation constant (K_d_) for LILRB1 ECD binding to RIFIN#1 was approximately 650 nM, which is consistent with the previously reported affinity determined using surface plasmon resonance (SPR). In contrast, the affinity of D1D2.v-IgG binding to RIFIN#1 was significantly higher, with a K_d_ of 6 nM, a 110-fold increase (Fig. 3c). To assess the ability of D1D2.v-IgG to antagonize the binding of LILRB1 to RIFIN#1, we performed a competitive ELISA assay. The results showed that D1D2.v-IgG effectively inhibited the binding of LILRB1 to RIFIN#1 in a dose-dependent manner with a half-maximal inhibitory concentration (IC_50_) of 20 nM (Fig. 3d).

**Fig 3:**
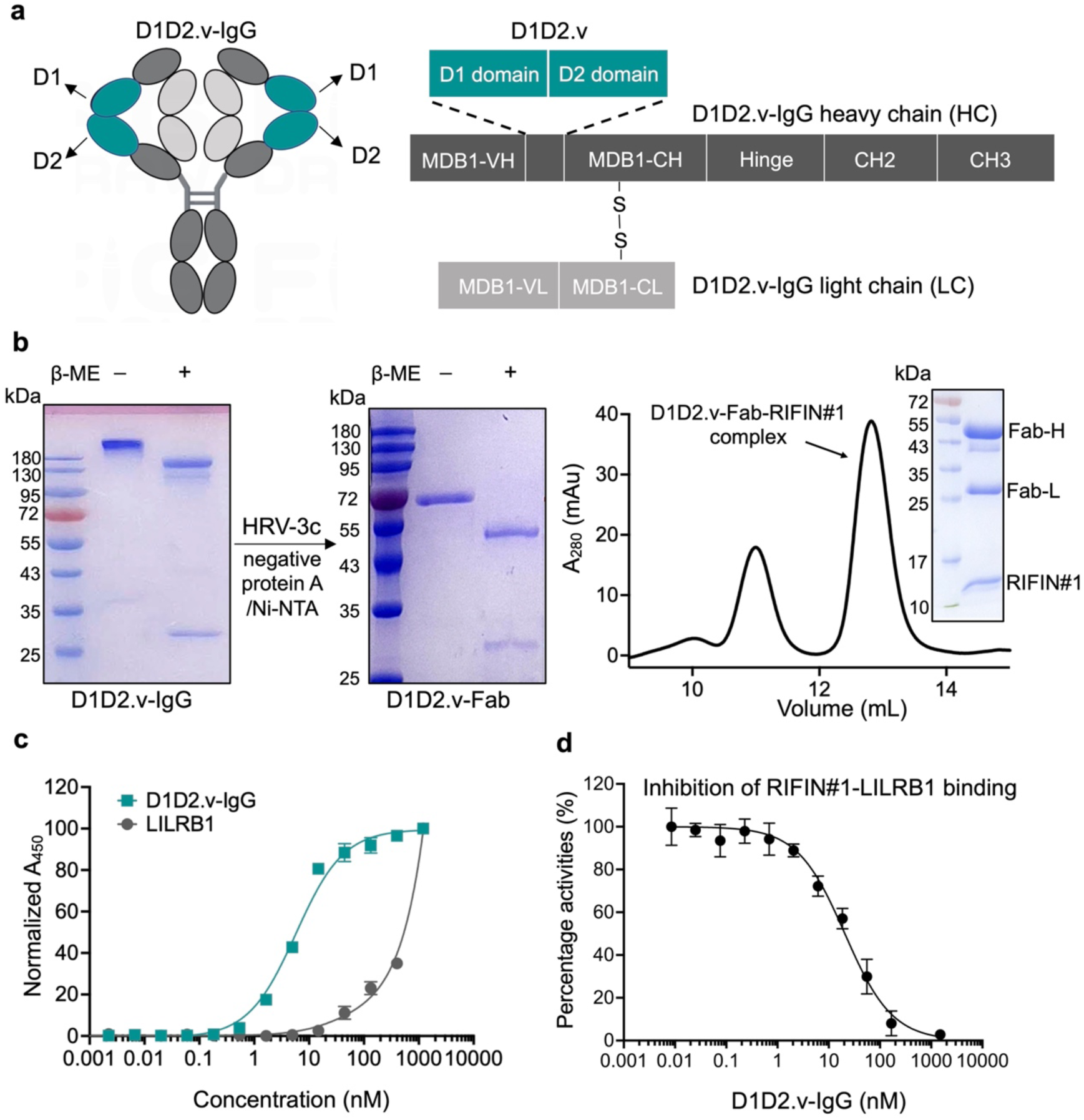
Molecular design and biochemical characterization of the D1D2.v-IgG. **(a)** Molecular structure and cDNA frame of LILRB1 D1D2-containing IgG. The inserted LILRB1-D1 domain is colored in dark blue and LILRB1-D2 domain is colored in light blue, which are located between heavy variable chain (VH) and heavy constant chain (CH). The antibody heavy chain and Fc domain are depicted in dark grey and the light chain is depicted in light grey. (**b)** Purification and functional validation of the antigen binding fragment (Fab) of D1D2.v-IgG. SDS-PAGE (Left) show the purity and molecular weight of D1D2.v-IgG before protease digestion. SDS-PAGE (Right) show the purity and molecular weight of D1D2.v-Fab. Size-exclusion chromatography (SEC) chromatograms show the homogeneous Fab-RIFIN complex have a UV_280_ absorbance at around 13 mL. **(c)** Titration curves of wild-type LILRB1 (ECD) (dark green) and D1D2.v-IgG (red) binding to RIFIN in ELISA assay. Data are mean ± SD. (**d)** The representative competition binding curve using wild type LILRB1 (ECD) show D1D2.v-IgG antagonizes the RIFIN-LILRB1 (ECD) interaction effectively. Data represent mean ± SD of three technical replicates.

### D1D2.v-IgG significantly enhances NK92 cytolytic activity against RIFIN#1-expressing K562 cells in vitro

Given that RIFIN#1 binding to LILRB1 inhibits NK cell responses, we subsequently investigated whether D1D2.v-IgG could enhance NK cell-mediated cytotoxicity by blocking the interaction between LILRB1 and RIFIN#1. To assess this, we performed NK92 cell cytotoxicity assays on K562 cells stably transfected with full length RIFIN#1 (Fig. S7). Flow cytometry verified the expression level of LILRB1 on NK92 cells (Fig. 4a). The RIFIN#1-expressing K562 cells were co-cultured with NK92 cells at two different effector to target (E: T) ratios of 2.5 and 12.5 in the presence or absence of D1D2.v-IgG. Flow cytometry analysis confirmed the specific binding of purified D1D2.v-IgG to the RIFIN#1-expresssing K562 cells (Fig. 4b). The cytotoxicity against K562 cells expressing RIFIN#1 was 5.8% in the absence of D1D2.v-IgG when the E:T ratio was 2.5, and increased significantly to 13.8% when D1D2.v-IgG at a concentration of 100 nM was added. Similarly, when the E:T ratio was 12.5, the cell killing efficiency was 35% in the absence of D1D2.v-IgG, but increased significantly to 61% when D1D2.v-IgG at a concentration of 100 nM was added (Fig. 4c). In addition, we observed that D1D2.v-IgG significantly increased the secretion of perforin and granzyme B from NK92 cells (Fig. 3d-e). These findings clearly indicate that D1D2.v-IgG can effectively stimulate NK-cell cytotoxicity against RIFIN#1-expressing K562 cells.

**Fig 4:**
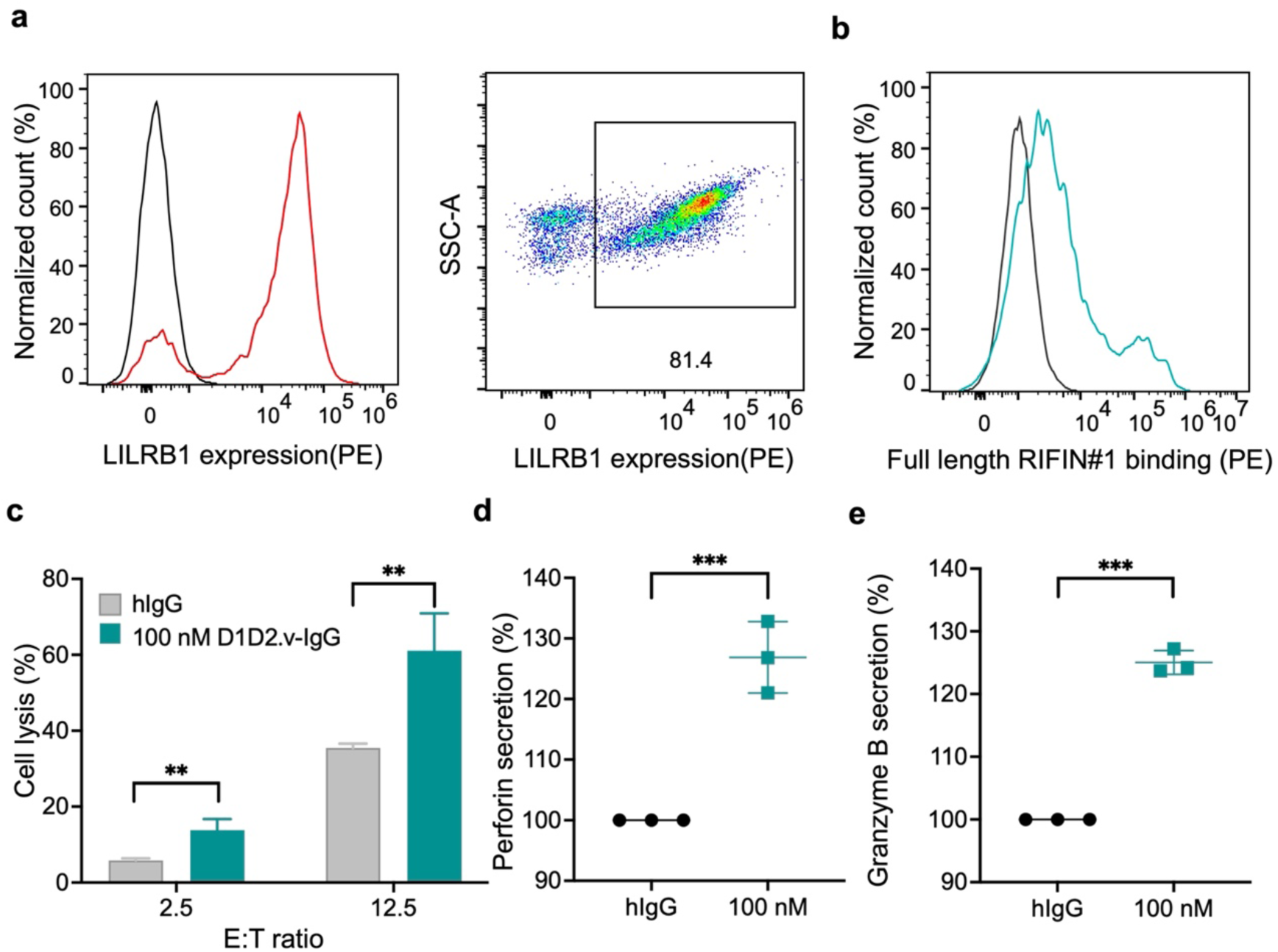
D1D2.v-IgG enhanced NK cell cytotoxicity against RIFIN-expressing K562 cell. (**a)** Detection of the LILRB1 on NK92 cells by flow cytometry. The red curve shows the binding of LILRB1 antibody to LILRB1 on NK92 cells, while the blue curve informs the control sample of NK92 cells. (**b)** Detection of the purified D1D2.v-IgG bound to RIFIN#1-expressing K562 cells by flow cytometry. The red curve shows the binding of D1D2.v-IgG to full-length RIFIN#1 on stably-transfected K562 cells, while the cyan curve informs the blank sample of K562 cells. **(c)** The results for the NK92 cytotoxicity in the presence or absence of D1D2.v-IgG. RIFIN#1-expressing K562 cells were cocultured with NK92 cells and incubated with 100 nM control human IgG (hIgG) (Grey, n = 3) or 100 nM D1D2.v-IgG (Yellow, n = 3). (**d,e)** Detections of the perforin (e) and granzyme B (f) releases in the presence or absence of D1D2.v-IgG. RIFIN#1-expressing K562 cells were cocultured with NK92 cells (E: T ratio = 1: 1) and incubated with 100 nM control hIgG (Grey, n = 3) or 100 nM D1D2.v-IgG (Orange, n = 3). After the incubation for 24h, the perforin and granzyme B levels were determined by ELISA. Statistical test was t test and statistical significance was analyzed between D1D2.v-IgG and hIgG. *P<0.05; **P<0.01; ***P<0.001; ****P<0.0001.

### Generation of a bispecific antibody targeting RIFIN#1 and NKG2D

Next, we developed a bispecific antibody that recruits NK cells to RIFIN#1-expressing cells using the D1D2.v-IgG as a basis. The redirection of NK cells requires a highly specific and high-affinity ‘compass’ (Fig. 5a). To fulfill this requirement, we chosen an engineered scFv called KYK2.0, which exhibits a nanomolar affinity towards NKG2D expressed on all types of NK cells. By fusing the scFv form of KYK2.0 to the C terminal of the Fc fragment of D1D2.v-IgG via a flexible GS linker, we generated a bispecific antibody (bsAb), named NK-biAb (Fig. 5b). The resulting bsAb was then transiently expressed and purified from Expi293F cell culture supernatants by affinity chromatography. Integrity and purity of NK-biAb were analyzed by SDS-PAGE under reducing and non-reducing conditions (Fig. S8a). As exprected, this purified bispecific antibody demonstrated the ability to bind to both NK cells and RIFIN#1-expressing K562 cells in flow cytometry analysis (Fig. S8b-c).

**Fig 5:**
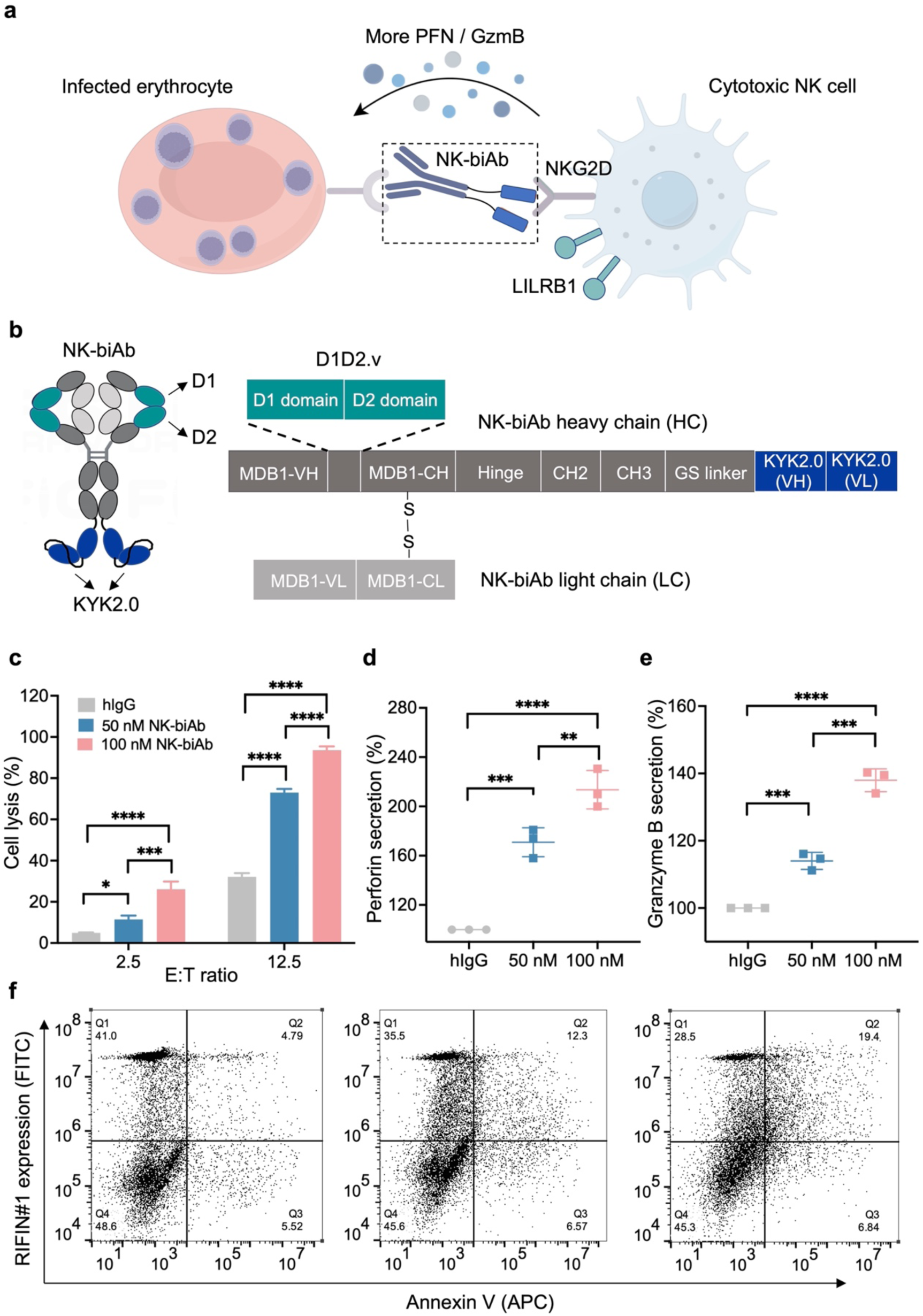
Design and characterization of NK-biAb. **(a)** Schematic of NK cell redirections by NK-biAb. Scfv form of KYK2.0 (scfv-KYK2.0), an NKG2D activating antibody was fused to C terminus of D1D2.v-IgG. When recognizing and blocking immune escape response of *P. falciparum* via D1D2.v-Fab domains, scfv-KYK2.0 domain can also bind to NKG2D receptors, redirect NK cells to *P. falciparum* and trigger NK cell activities. scFv-KYK2.0 is colored in light orange and NKG2D receptor is depicted as a light blue dart. **(b)** cDNA frame of NK-biAb. cDNA of D1D2.v-IgG part is the same as previously described. The heavy variable chain (VH) and light variable chain (VL) of KYK2.0 depicted in orange are fused to C terminus of D1D2.v-IgG by a GS linker. (**c)** NK92 cytotoxicity assay in the presence of NK-biAb. RIFIN#1-expressing K562 cells were cocultured with NK92 cells and incubated with 100 nM control human IgG (hIgG) or 50 nM NK-biAb (Cyan, n =3) and 100 nM NK-biAb (Red, n =3). Statistical tests were two-way ANOVA and statistical significance was analyzed between NK-biAb and hIgG. **(d,e)** Detections of the perforin (d) and granzyme B (e) releases in the presence or absence of NK-biAb. RIFIN#1-expressing K562 cells were cocultured with NK92 cells (E: T ratio = 1: 1) and incubated with 100 nM control hIgG (Grey, n = 3), 50 nM NK-biAb (Cyan, n =3) and 100 nM NK-biAb (Red, n = 3). After the incubation for 24h, the perforin and granzyme B levels were determined by ELISA. Statistical tests were one-way ANOVA and statistical significance was analyzed between NK-biAb and hIgG. **(f)** Flow detection results for the coculture of NK92 and RIFIN#1-expressing K562 for 24 h at E: T = 1: 1 in the presence of 100 nM hIgG (Left), 50 nM NK-biAb (Middle) and 100 nM NK-biAb (Right), showing the RIFIN#1-expressing K562 cells killed by NK92 cells. The data in the second quadrant stands for the lysed RIFIN#1-expressing K562 cells. *P<0.05; **P<0.01; ***P<0.001; ****P<0.0001.

### NK-biAb exhibits an enhanced ability to trigger NK92 cytotoxicity against RIFIN#1-expressing K562 cells

Finally, we evaluated the effect of NK-biAb on NK cell cytotoxicity against RIFIN#1-expressing K562 cells. As shown in figure, the ability of this bispecific antibody to trigger NK cytotoxicity was significantly enhanced compared to the monospecific antibody D1D2.v-IgG. When treatment with NK-biAb at a concentration of 100 nM, there was a remarkable enhancement in cytotoxic efficiency from 13.8% to 26.1% and from 61%-93.6%, at E/T ratios of 2.5 and 12.5 respectively, compared to treatment with D1D2.v-IgG under the same conditions (Fig. 5c). Compared with D1D2.v-IgG treatment, NK92 cells secreted more perforin and granzyme B in the presence of 50 nM and 100 nM NK-biAb (Fig. 5d-5e). Flow cytometry results also verified that NK-biAb can potently stimulate NK-cell cytotoxicity against RIFIN#1-expressing K562 cells (Fig. 5f).

## Discussion

Upon infection, *P. falciparum* utilizes variant surface antigens to engage with receptors on the human host, thereby facilitating immune evasion and ensuring their survival in the host^7,8^. Recent studies have identified several receptor-containing antibodies in malaria-exposed individuals that block surface antigen-receptor interactions. Functional studies have shown that these antibodies hold potential as therapeutic agents and for incorporation into malaria vaccine design^15–17^. For instance, MGD21 antibody has been shown to promote NK cell-dependent cleavage of IEs and to inhibit the growth of *P. falciparum*^17^. However, reported receptor-containing antibodies only have either LAIR1 domain or LILRB1-D3D4 domains, and no antibodies containing LILRB1-D1D2 domains have been discovered. The RIFIN family is divided into different species, some of which interact with LILRB1-D1D2 domains rather than LILRB1-D3D4 domain or LAIR1 to transduce immunoinhibitory signals. Therefore, the use of LILRB1 D1D2-containing antibodies for the treatment of malaria is of great importance.

In this study, we propose an in vitro strategy for designing novel receptor-containing antibodies. Our approach utilizes structure-guided affinity maturation to obtain receptor fragment variants with enhanced binding capacity, while using the framework of a human natural antibody against malaria as a template for antibody generation^15^. Due to the weak affinity between RIFIN and LILRB1-D1D2, no relevant D1D2-containing antibody has been reported. We used this strategy to artificially create high-affinity D1D2-containing antibodies, filling a gap in this field. As an illustrative example, we successfully prepared D1D2.v-IgG antibody, which has 110-fold higher affinity towards RIFIN1# than LILRB1 and can effectively block the interaction between RIFIN1# and LILRB1. Cellular experiments demonstrated that D1D2.v-IgG significantly enhanced NK cell-mediated cytotoxicity against K562 cells expressing RIFIN#1. In addition, on the basis of D1D2.v-IgG, we developed a bispecific antibody targeting both RIFIN#1 and the NKG2D receptor, which showed enhanced ability to trigger NK cell-mediated cytotoxicity compared to D1D2.v-IgG. Bispecific antibodies have been shown to be a promising avenue for immunotherapeutic interventions in a variety of malignancies^21,22,24,26^. Our data emphasize the potential application of these agents in combating malaria.

In conclusion, our study provides a strategy for the design of receptor-containing antibodies, offering potential applications in the development of malaria therapeutics and vaccines. The increasing availability of resolved structures of Plasmodium surface antigens binding to host receptors^9,11,15,17^ will facilitates the development of corresponding antibodies using our strategy.

## Materials and Methods

### Antigen purifications

The variable region of RIFIN#1 (PF3D7_1254800, 165-274 aa) with an N-terminal 6×His tag was cloned into a pET28a vector. The recombinant vector was transformed into E.coli BL21 (gold), and the cells were induced with 1 mM IPTG at a OD_600_ of 0.7. Protein was expressed as inclusion bodies and denatured in a guanidine buffer (6 M Guanidine hydrochloride, 20 mM Tris pH 8.0, and 300 mM NaCl) for 4 h. The inclusion bodies were then purified by Ni-NTA affinity chromatography and eluted by the guanidine buffer containing additional 500 mM imidazole. The denatured inclusion bodied were retrieved by dialyzing against a buffer (20 mM Tris pH 8.0, 300 mM NaCl) and resolubilized in the guanidine buffer. Refolding was performed by the quick dilution of inclusion bodies into a buffer (1 M L-arginine, 20 mM Tris pH 8.5, 100 mM NaH_2_PO_4_, 2 mM ethylenediaminetetraacetic acid, 0.1 mM phenylmethylsulfonyl fluoride, 3 mM reduced glutathione and 0.3 mM oxidized glutathione) until a final protein concentration of 0.1 mg/mL-0.2 mg/mL. The refold RIFIN#1 was further polished on a Superdex 75 Increase column in a PBS (pH 7.2) buffer. The ECD of human LILRB1 gene with a C-terminus His tag was cloned into a pTT5 vector, and the vector was then PEI-transfected into Expi293F cells (Procell, China). After a post-transfection for 96 h, the supernatant was collected and the protein was purified with a Ni-NTA affinity chromatography. The purified LILRB1-ECD was further exchanged into PBS (pH 7.2) buffer on a Superdex 200 Increase column.

For biotinylation, the RIFIN#1 and LILRB1-ECD proteins were cultured with NHS-pEG12-biotin in a PBS buffer (pH 7.2). The culture reactions were quenched with a Tris buffer after 8 h incubation. The free NHS-pEG12-biotin reagents were splitted from the biotin-labeled proteins by a size-exclusion chromatography.

### EBY100 vector construction and LILRB1-D1D2 wild type display

To avoid weird display stuff, pYD1 vector was reconstructed by fusing Agar 1/2 domain to the C-terminus of displayed targets. Meanwhile, the detection tags of pYD1 vector were replaced by a N-terminal FLAG, a C-terminal V5-tag and a C-terminal His tag. D1D2 wild type was co-electroporated with a precut reconstructed pYD1, while yeast cells were induced with SGCAA. The signals of RIFIN#1 binding to the displayed D1D2 wild type were detected by FCM.

### Constructing the library of D1D2 variants

Based on the crystal structure of RIFIN#1-LILRB1 complex, a site-directed library was generated by the several rounds of overlapping PCR reactions for randomizing the binding residues of LILRB1-D1D2. The randomizing primers are as shown in Supplementary Table 1.The overlapping PCR reactions were conducted under 20 cycles. The external primers for the amplification of LILRB1-D1D2 library on a large scale introduce a 70 bp homologous sequence to pYD1 vector. The LILRB1-D1D2 library were co-transformed into EBY100 with a precut pYD1 vector at a library/vector ratio of 3: 1. Yeast transformants were recovered in a YPD/Sorbitol broth and amplified in SDCAA, resulting in a library containing 2×10^7^ transformants.

### MACS selections of the naive library

Transformed yeast were induced in SGCAA medium at 20°C for 48 h. For the 1^st^ round selection, 4×10^8^ cells referred to as 20× library size were pellet and washed with a PBE buffer (PBS pH 7.2 supplemented with 0.5% BSA). Firstly, the cells were resuspended in 2 mL of the PBE buffer containing 2 µM biotinylated RIFIN that had been pre-coated with Streptavidin-Alexa Fluor 647 (Invitrogen, USA), followed by an incubation at 4°C for 2 h. Secondly, the stained cells were pelleted by centrifugation (3500 g, 5 min), washed twice with 2 mL PBE, and resuspended in PBE containing paramagnetic anti-Alexa Fluor 647 microbeads (Miltenyi Biotech, German). Thirdly, a magnetically-labeled yeast cells were enriched on LS MACS column (Miltenyi Biotech, German), and the enriched yeast cells were eluted with 5-10 mL PBE. Finally, the selection outputs were pellet and amplified in SDCAA for next round selection. Five additional selections were performed by successively decreasing concentrations of biotinylated RIFIN#1 from 1 µM to 0.02 µM. In each round selection, the yeast cells were first labeled with SAV-Alexa Fluor 647 after incubating with biotinylated RIFIN#1 for 2 h and washed once with PBE, then coated with 100 µL of anti-Alexa Fluor 647 microbeads for 30 min and washed once, and finally separated by LS MACS column.

### MACS competition selections

The MACS tetramer/monomer selection outputs were further performed competition selections to get ultrahigh affinity variants. The induced tetramer/monomer selection outputs were cultured with 0.02 µM biotinylated RIFIN#1 at 4°C for 2 h. Then the cells were pellet, washed once with PBE and competed with 1 µM unlabeled RIFIN#1 for 2 h. After the competition process, the cells were stained with Streptavidin-Alexa Fluor 647 and coated with anti-Alexa Fluor 647 microbeads. MACS sorting was also performed using LS column. By now, the 1^st^ round competition was finished, and the competition outputs were amplified and induced for next round competition selection. In the 2^nd^ round competition selection, the competing time of unlabeled RIFIN was increased to 4 h, and the final selection outputs were titrated on SDCAA plates for plasmid extraction and sequencing.

### Preparation of D1D2.v-IgG

D1D2.v were inserted between MDB1-VH and MDB1-CH, and cloned into pTT5 vector as IgG4 forms. MDB1 light chain was also cloned to pTT5 vector. The DNAs of MDB1 light chains and D1D2.v-IgG heavy chains were co-transfected into Expi293F cells using PEI transfection reagents. After the transfection for 96 h, the culture supernatant was separated by centrifugation (4000 rpm, 10 min) and purified with protein A resin. The purified D1D2.v-IgG was further polished on a Superdex 200 Increase column in a PBS (pH 7.4) buffer.

### Characterization of D1D2.v-IgG

Protein-based ELISA assays were performed to determine the affinity of D1D2.v-IgG to RIFIN#1 in comparison to that of LILRB1-ECD. First, RIFIN#1 (5 µg/mL) was immobilized on 96-well ELISA plates and incubated with D1D2.v-IgG of different concentrations. Then, the plates were washed with PBST for 3 times and cultured with the anti-human IgG-HRP at 1: 5000 dilution. Finally, the wells were washed with PBST, developed with TMB for 15 min, and quenched with 2 M HCl. The absorbance at 450 nm was determined by a microplate reader.

For the protein-competition ELISA assay, RIFIN#1 (5 µg/mL) was immobilized on 96-well ELISA plates, and cocultured with D1D2.v-IgG of different concentrations and a competing biotinylated LILRB1 (ECD) of 1-1.5× *K*d concentration, where *K*_d_ is the dissociation constant. Then the plates were washed with PBST for 3 times and cultured with HRP-conjugated streptavidin at 1: 2000 dilution. The absorbance at 450 nm was recorded by a microplate reader.

### Purifying Fab fragment of D1D2.v-IgG and its functional validation

D1D2.v-IgG was digested by His-tagged HRV-3c protease overnight and subjected to a negative protein A and Ni-NTA column purification. The 5-fold excess of refold RIFIN#1 were co-cultured with the Fab fragment of D1D2.v-IgG and the resulting complex was further polished on a Superdex 200 Increase column. The ratio of RIFIN#1 binding to the Fab fragment of D1D2.v-IgG was assessed by SDS-PAGE.

### Preparation of NK-biAb and its functional validation

scFv-KYK2.0 was fused to the C-terminus of D1D2.v-IgG heavy chain to generate NK-biAb (*M*_r_ = 260 kDa). The DNAs of MDB1 light chains and NK-biAb heavy chains were co-transfected into Expi293F cells using PEI transfection reagents. After the transfection for 96 h, the culture supernatant was collected by centrifugation (4000 rpm, 10 min) and purified on a protein A column. The purified protein was further polished on a Superdex 200 Increase column in a PBS (pH 7.4) buffer. To validate the functions of NK-biAb to attach NK92 cells and bind NKG2D, the NK92 cells pre-stimulated with IL-2 were cultured with NK-biAb at 37°C for 2 h, and stained with anti-human IgG-PE for the detection of surface NKG2D.

### Stable RIFIN#1-expressing K562 cell line production

K562 cells (Procell, China) were cultured at 37°C in DMEM with 10% FBS and 5% CO_2_. The cDNA of RIFIN#1 with its natural signal peptide was cloned into pLVX-ZsGreen plasmid. Lentivirus were produced by HEK-293T (Procell, China) for infecting K562 cells. The infected K562 cells were enriched by the selection using 1 μg/mL puromycin, to obtain the stable RIFIN#1-expressing K562 cell line.

### Evaluation of NK cell cytotoxicity

The RIFIN#1-expressing K562 cells were incubated at 37°C for 1 h with D1D2.v-IgG or NK-biAb. The NK92 cells in the same volume as RIFIN#1-expressing K562 cells were then cocultured for 4 h at 37°C and 5% CO_2_ with the incubated RIFIN#1-expressing K562 cells at the E: T ratios of 2.5: 1 and 12.5: 1. The final mixtures contain either 100 nM D1D2.v-IgG or 50 nM/100 nM NK-biAb. The killing efficiency (*E*) was calculated from the absorbance (*A*) of CCK8 at 450 nm: *E*% = [1−(*A*_S_−*A*_EC_)/*A*_TC_]×100%, where *A*_S_ is for the mixture of effector and target cells, and *A*_EC_ and *A*_TC_ are, respectively, for effector cells and target cells.

### Cytokine measurement and FCM detection

10^5^ NK92 cells were cocultured for 24 h with 10^5^ RIFIN#1-expressing K562 cells in a 96-well plate in the presence of either 100 nM D1D2.v-IgG or 50 nM/100 nM NK-biAb. Perforin and granzyme B release were detected from the supernatants of the cocultured mixture by protein-based ELISA assay kits (Beyotime, China). The mixtures of NK92 and K562-RIFIN#1 cells were stained for 1 h at 4°C (controlled by ice) with Annexin V-APC in 100 μL PBS containing 0.5% BSA. The stained cells were detected by flow cytometry, with the data analyzed by Flowjo software.

## Acknowledgements

Funding was provided by the National Key R&D Program of China (Grant No. 2023YFA1607500), the National Natural Science Foundation of China (Grant No. 32371362), and the Anhui Major Basic Research Projects (Grant No. 2023z04020016).

## Author contributions

Y.W and H.T contributed equally to this work. Y.W initiated the project, performed the library construction and selections, prepared antigen samples and performed biochemical assays. H.T performed the FCM detections and conducted the stable cell line production, cytotoxicity assays and ELISA assays. W.W performed the antibody purifications and functional validations. L.M helped to prepare biotinylated sample preparations. C.Z helped to perform biochemical assays and FCM detections. Y.W. and B.W analyzed the data, interpreted the results. J.W, H.D and B.W prepared the manuscript. J.W., B.W and H.Z supervised the project. All authors read and edited the manuscript.

## Data and materials availability

All data and materials are available within the paper and its supplementary materials. Extra data are available from the corresponding author (J. Wang) upon reasonable request.

