## supplemental information for "Potent LILRB1 D1D2-containing antibodies inhibit RIFIN-mediated immune evasions"

**Supplementary Figures**

**b**

**Fig. S1**

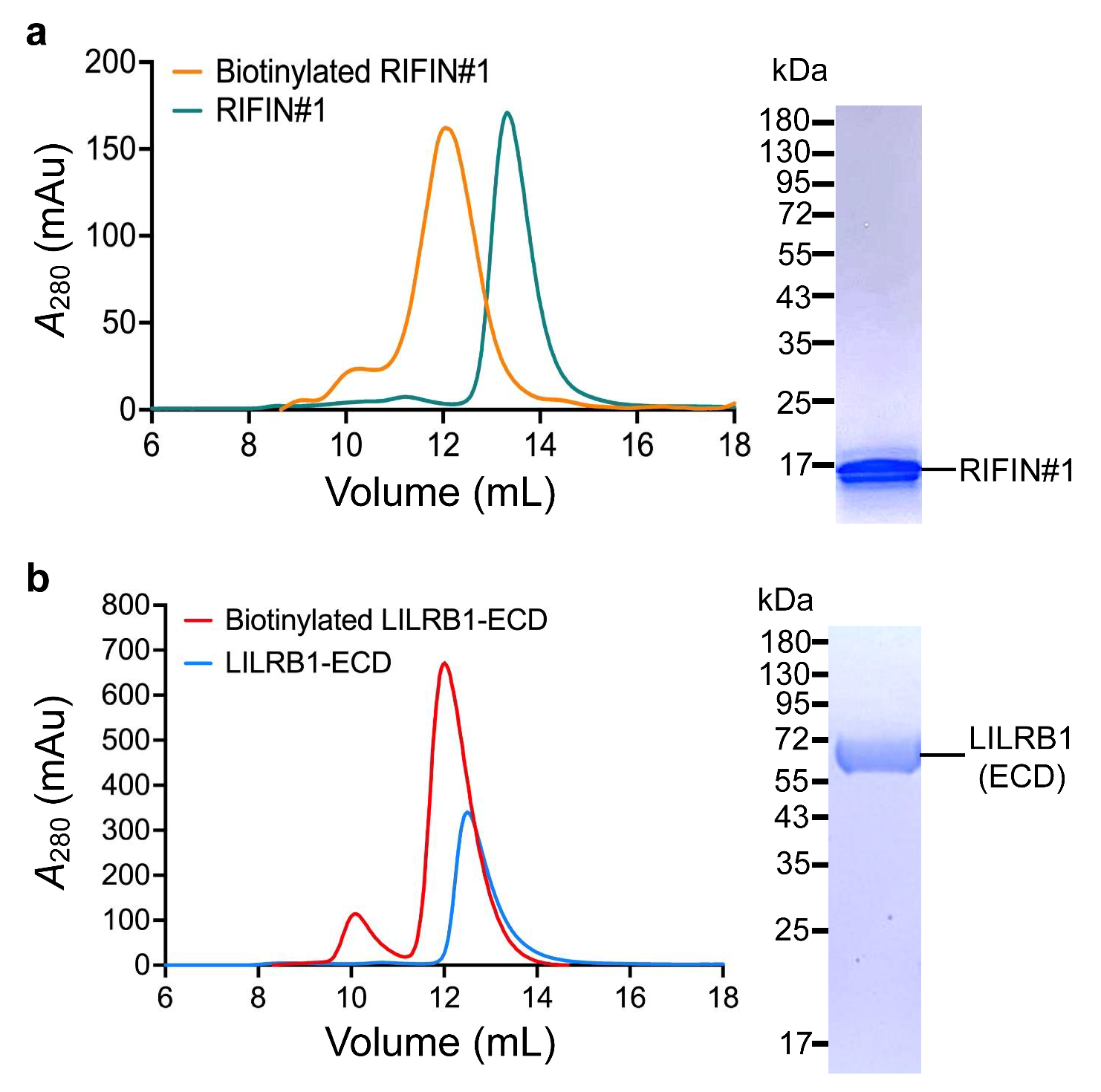

**Fig. S1: Protein purifications and labeling.** **(a)** Size-exclusion chromatography chromatograms from the refolding and purification of biotinylated and non-biotinylated RIFIN#1. RIFIN#1 was purified by nickel affinity chromatography and SDS–PAGE gels showing the purity and molecular weight. **(b)** Size-exclusion chromatography chromatograms from the purification of biotinylated and non-biotinylated LILRB1 (ECD). LILRB1 (ECD) was purified by nickel affinity chromatography and SDS–PAGE gels showing the purity and molecular weight**.**

**Fig. S2**

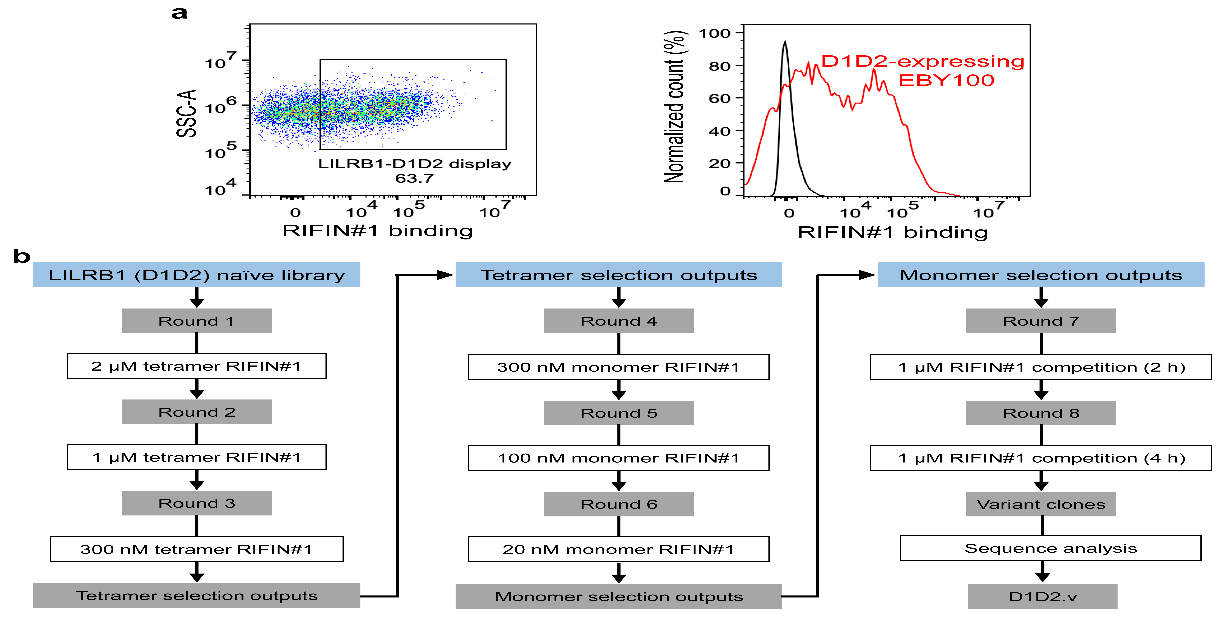

**Fig. S2: Correct folding of displayed LILRB1-D1D2 domains and schematic of D1D2 variant selections.** **(a)** Functional folding of LILRB1-D1D2 domains on EBY100 surface. The RIFIN#1 binding signal indicates the correct structure of LILRB1-D1D2 domains on EBY100 surface. **(b)** Flow chart depicting the selection strategy used to isolate high-affinity D1D2 variants (D1D2.v). Three different selection strategies were performed until the high affinity D1D2 variant sequences were conserved.

**Fig. S3**

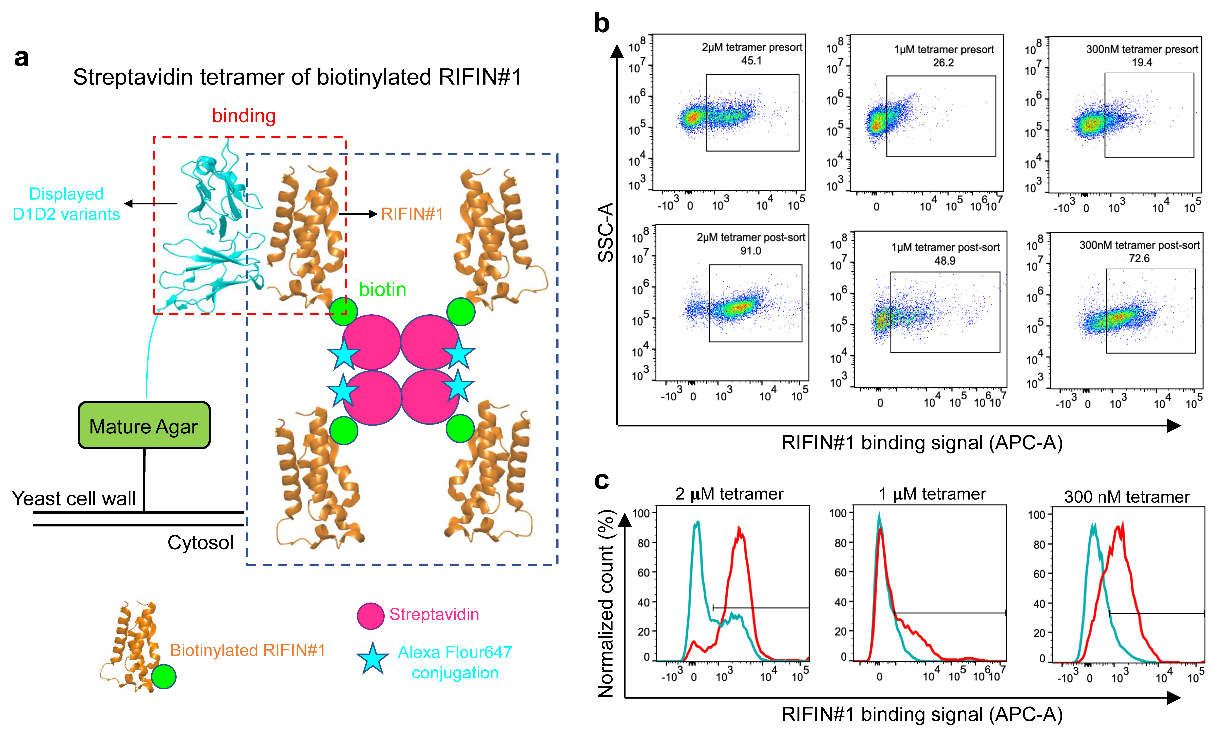

**Fig. S3: Library selections using RIFIN#1-tetramer.** **(a)** Schematic of tetramer selection. Three rounds of selections were performed for retaining D1D2 variants bound to the RIFIN#1-tetramer to remove non-binders or low-affinity binders towards RIFIN#1. **(b)** The positive binding rate of yeast cells towards RIFIN#1-tetramer before and after each round of tetramer selection. **(c)** The validity of each round of tetramer selection. The Presort library are coloured in cyan and post-sorting outputs were coloured in red.

**Fig. S4**

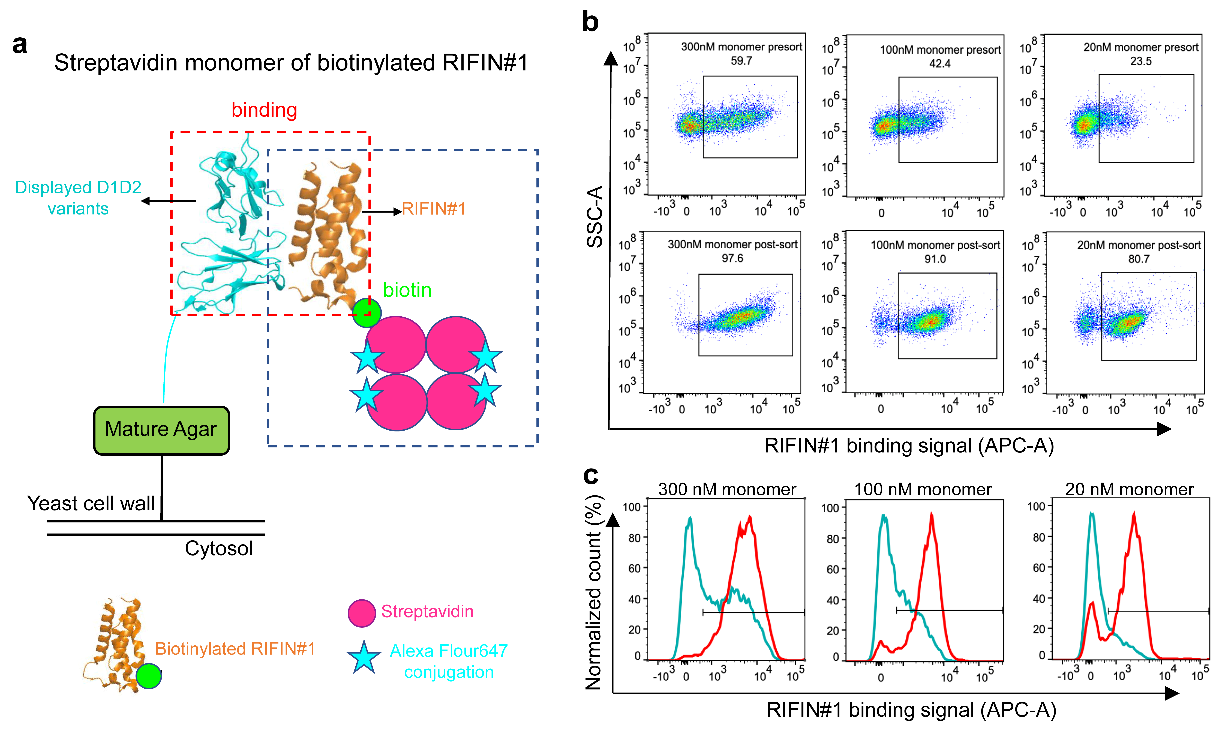

**Fig. S4: Library selections using RIFIN#1-monomer. (a)** Schematic of monomer selection. Three rounds of selections were performed for enriching RIFIN#1-binders with mid-to-high affinity. **(b)** The positive binding rate of yeast cells towards RIFIN#1-monomer before and after each round of monomer selection. **(c)** The validity of each round of monomer selection. The presort library are coloured in cyan and post-sorting outputs were coloured in red.

**Fig. S5**

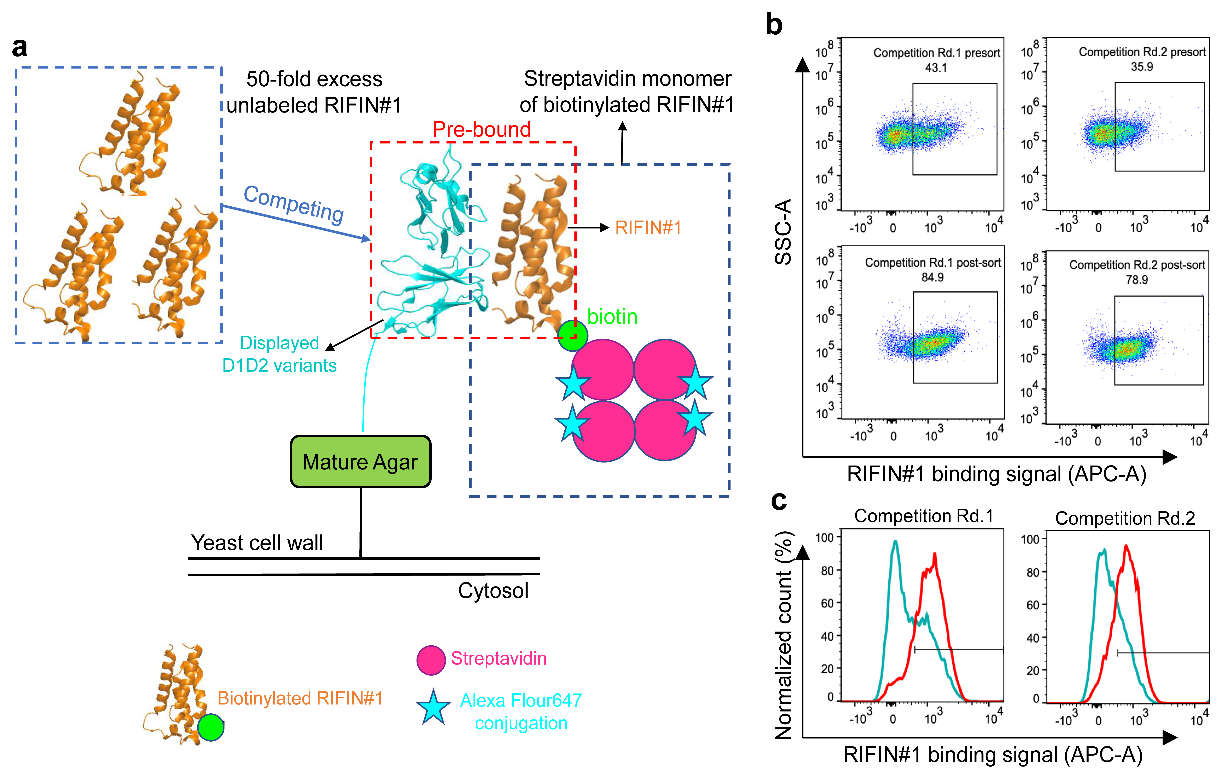

**Fig. S5: Competition selections. (a)** Schematic of competition selection. Two rounds of competition selections were performed for further enriching the high affinity RIFIN#1-binders, where 50-fold excess unbiotinylated RIFIN#1 was used to compete with prebound biotinylated RIFIN#1-monomer for 2h and 4h respectively. **(b)** The positive binding rate of yeast cells towards RIFIN#1-monomer before and after each round of competition selection. **(c)** The validity of each round of competition selection. The presort library are coloured in cyan and post-sorting outputs were coloured in red. The diversity of D1D2 variants tend to be conserved.

**Fig. S6**

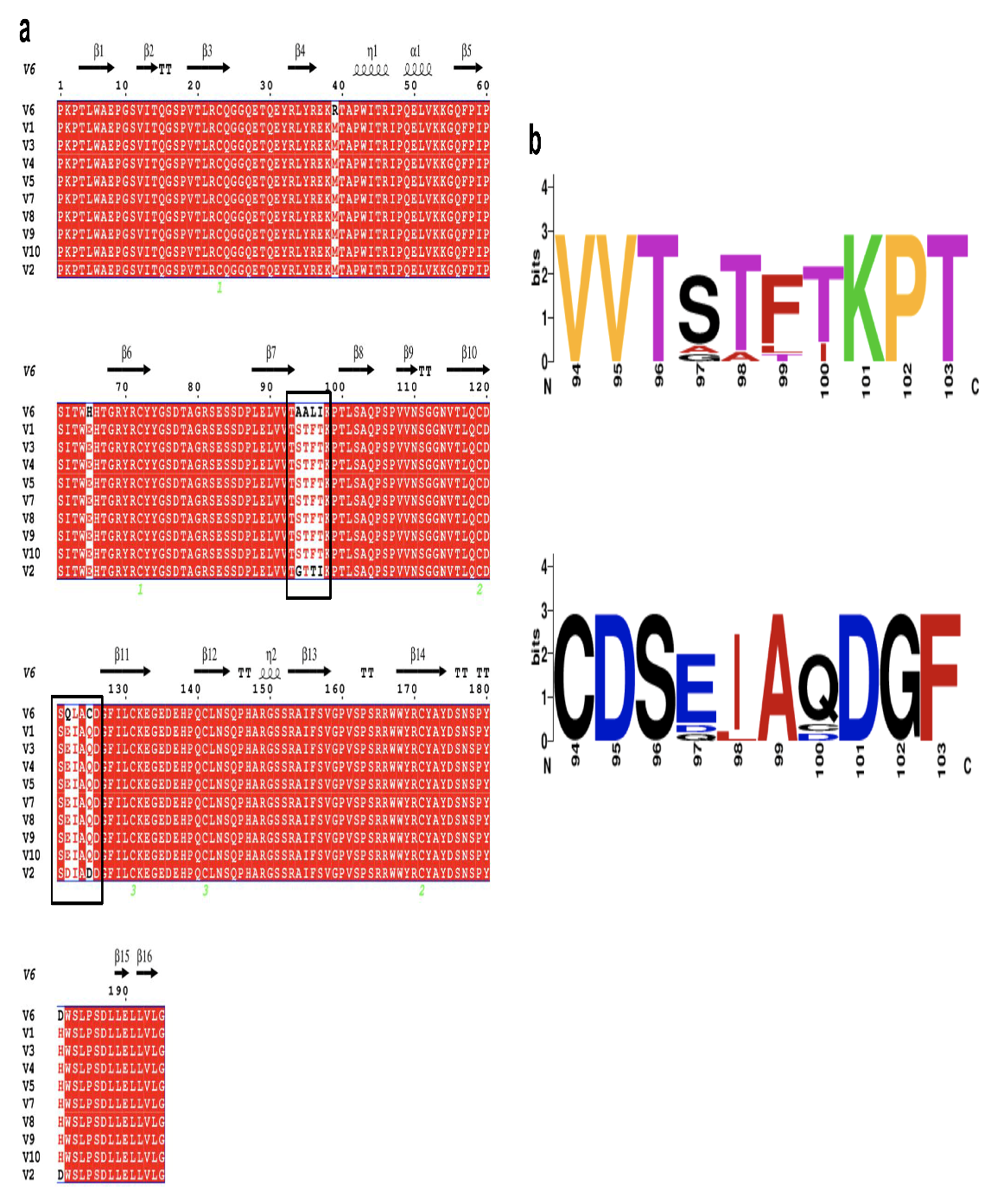

**Fig. S6: Sequence alignment of yeast clones after eight-round of selections. (a)**Ten random yeast clones were sequenced and sequence alignment was performed on Clustal Omega. The sequence difference was displayed by ESPript3.0. Residues that contact with RIFIN#1 are coloured in white and the main binding cores are framed in black. 80% of these ten clones contain a same sequence D1D2.v. **(b)** Sequence logos for the residues of the main binding cores of LILRB1-D1D2 variants.

**Fig. S7**

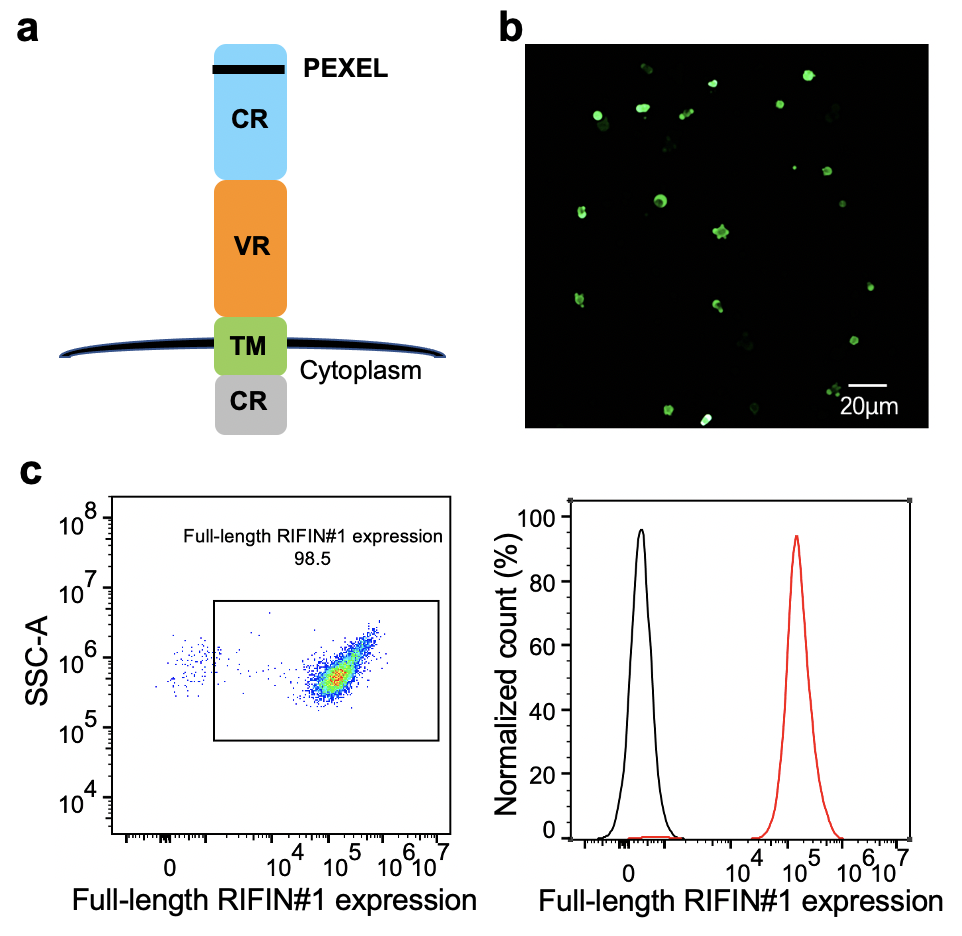

**Fig. S7: RIFIN#1 construct and full-length RIFIN expression. (a)** Expression construct of wild type full length RIFIN#1 on K562 cell membrane. **(b)** Detection the morphology of K562 cells stablely expressing full-length RIFIN#1. **(c)** Measurement of expression rate of K562 cells stablely expressing full-length RIFIN#1 by flow cytometry. The rate of K562 cells expressing full-length RIFIN#1(Red) is 98.5% compared to blank K562 cells (Black), which indicates full-length RIFIN#1 causes low toxicity to K562 cells and K562-RIFIN#1 cell line exists stably.

**Fig. S8**

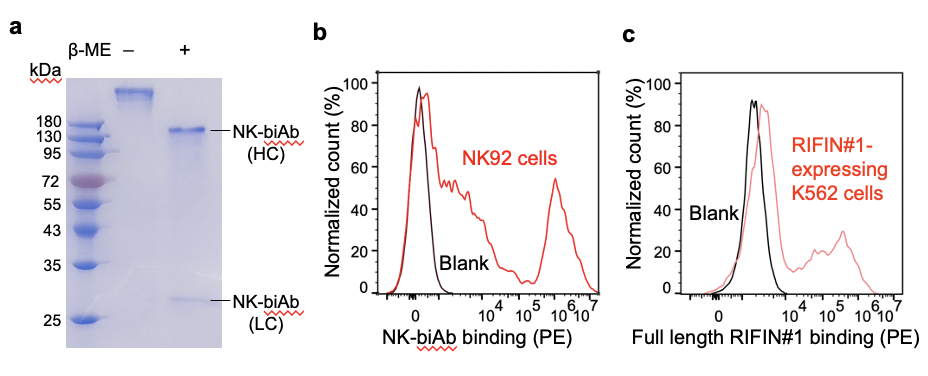

**Fig. S8: NK-biAb purification and functional validation. (a)** Protein purification of NK-biAb. Results were assessed by SDS-PAGE. **(b)** Functional validation of NK-biAb binding to the NKG2D on NK92 cells (Red), compared to blank sample (Black). **(c)** Functional validation of NK-biAb binding to the RIFIN#1 on K562 cells (Red), compared to blank sample (Cyan).

**Supplementary Table**

**Table 1. Library construction primers and wobble codons**

| **Primer** | **Sequence** |
| --- | --- |
| lib_K42-R | GGCCCTTCTTCACAAGCTCCTGTGGGATCCGTGTAATCCAGGGTGCTGTCNTCTTTTCT |
| lib_E68-F | AGGAGCTTGTGAAGAAGGGCCAGTTCCCCATCCCATCCATCACCTGGSAWCACA |
| lib_97-100-R | CCTGAGTTCACCACGGGGCTGGGCTGGGCTGAGAGGGTGGGTTTARWMNNASYASYTGTCACCACC |
| lib_125-128-F | AGCCCCGTGGTGAACTCAGGAGGGAATGTAACCCTCCAGTGTGACTCASAWNTTGCANNKGATGGCTTCATTC |
| lib_E184-R | AAGGGCCCTCTAGACTCGAGACCTAGGACCAGGAGCTCCAGGAGATCACTGGGTAGAGACCAWTSATAGGG |

| **Codon** | **Direction** | **residue** |
| --- | --- | --- |
| CNT | 3’—>5’ | K, R, T, M |
| SAW | 5’—>3’ | E, Q, D, H |
| ARW | 3’—>5’ | I, F, T, S |
| MNN | 3’—>5’ | All amino acids |
| ASY | 3’—>5’ | G, A, S, T |
| NTT | 5’—>3’ | F, I, L, V |
| NNK | 5’—>3’ | All amino acids |
| WTS | 3’—>5’ | E, Q, D, H |

Primers for LILRB1-D1D2 variant library constructions and wobble codons for randomizing LILRB1-D1D2.
